# Systematic classification error profoundly impacts inference in high-depth Whole Genome Shotgun Sequencing datasets

**DOI:** 10.1101/2022.04.04.487034

**Authors:** James Johnson, Shan Sun, Anthony A. Fodor

## Abstract

There is little consensus in the literature as to which approach for classification of Whole Genome Shotgun (WGS) sequences is best. In this paper, we examine two of the most popular algorithms, Kraken2 and Metaphlan2 utilizing four publicly available datasets. As expected from previous literature, we found that Kraken2 reports more overall taxa while Metaphlan2 reports fewer taxa while classifying fewer overall reads. To our surprise, however, Kraken 2 reported not only more taxa but many more taxa that were significantly associated with metadata. This implies that either Kraken2 is more sensitive to taxa that are biologically relevant and are simply missed by Metaphlan2, or that Kraken2’s classification errors are generated in such a way to impact inference. To discriminate between these two possibilities, we compared Spearman correlations coefficients of each taxa against each taxa with higher abundance from the same dataset. We found that Kraken2, but not Metaphlan2, showed a consistent pattern of classifying low abundance taxa that generated high correlation coefficients with higher abundance taxa. Neither Metaphlan2, nor 16S sequences that were available for two of our four datasets, showed this pattern. Simple simulations based on a variable Poisson error rate sampled from the uniform distribution with an average error rate of 0.0005 showed strikingly strong concordance with the observed correlation patterns from Kraken2. Our results suggest that Kraken2 consistently misclassifies high abundance taxa into the same erroneous low abundance taxa creating “phantom” taxa have a similar pattern of inference as the high abundance source. Because of the large sequencing depths of modern WGS cohorts, these “phantom” taxa will appear statistically significant in statistical models even with a low overall rate of classification error from Kraken. Our simulations suggest that this can occur with average error rates as low as 1 in 2,000 reads. These data suggest a novel metric for evaluating classifier accuracy and suggest that the pattern of classification errors should be considered in addition to overall classification error rate since consistent classification errors have a more profound impact on inference compared to classification errors that do not always result in assignment to the same erroneous taxa. This work highlights fundamental questions on how classifiers function and interact with large sequencing depth and statistical models that still need to be resolved for WGS, especially if correlation coefficients between taxa are to be used to build covariance networks. Our work also suggests that despite its limitations, 16S rRNA sequencing may still be useful as neither of the two most popular 16S classifiers showed these patterns of inflated correlation coefficients between taxa.

## Introduction

Improvements in sequencing technology have allowed researchers to explore novel relationship between microbes and their environments [1–4]. In tandem with the growth of 16S sequencing, two of the most popular classification algorithms in terms of past citations include Quantitative Insights into Microbial Ecology (QIIME) [5], which is often used for creation of closed-reference or open-referenced OTUs, and the Ribosomal Database Project (RDP) [6] which represents naïve Bayesian classification. While 16S sequencing remains powerful, it has several limitations: it can only reliably generate taxonomy at genus and above, functional information can only be inferred from taxonomy, it targets only a single gene, and the process of amplifying reads via Polymerase Chain Reaction (PCR) can introduce chimeras and other errors. Therefore, researchers have increasingly been incorporating WGS sequencing into microbiome analysis since it provides functional information about potential proteins, and can at least in theory, identify taxa down to the strain levels while tracking horizontal gene transfer events [7]. Unlike 16S sequencing, WGS sequencing amplifies entire genomes and it remains an open question how best to taxonomically classify WGS sequences.

Many algorithms have been proposed to address the research challenges associated with taxonomic classification of WGS sequences [8–19]. Attempts to benchmark these algorithms have led to conflicting results [20–24]. One paper by Peabody et al explored the precision and accuracy of WGS classifiers but argued that performance varied widely between different classifiers and advocated for the use of standardized datasets. In 2016, Lindgreen et al examined datasets they argued were more complex and more realistic than previous comparisons and found a high degree of variability between different tools. In 2017, McIntyree et al found that Kraken was among the tools with the highest read-level precision. In 2019, Ye et al argued that explicit consideration of reference databases should be an important consideration in evaluating and improving classifier performance. More recently in 2021, Sun et al argued that explicit consideration of sequence abundance vs. taxonomic abundance and how reference genome size was incorporated into normalization schemes is a crucial and underappreciated aspect of classifier performance.

All of the benchmarking papers referenced make extensive utilization of **in silico mock** (simulated mock) communities which comprise of samples created from random combinations of assembled sequences from either databases or previously sequenced datasets. These datasets have the obvious advantage that the “correct” answer is known but the obvious disadvantage that the mock communities might not resemble “real”, or naturally derived communities, in community structure and distribution. Moreover, choices made in assembling the mock communities can bias benchmarking results in favor of one or another class of algorithms. Similar concerns can be raised about **in vitro mock** communities which consist of known microbes with sequenced genomes grown in culture in the laboratory and mixed together in know proportions. In vitro mock communities are in some sense more “real” than in silico mock communities as they involve sequencing of actual microbes, but it is still unclear how choices made in assembling the mock community could influence benchmark results or how in vitro mock communities might miss important features of complex “real” communities. Ideally, benchmarking papers could utilize **naturally derived communities**, or “real” datasets, that consists of sequenced samples taken from the field or natural habitat of the microbiome under study with sample composition unknown prior to sequencing. While “real” samples are appropriately complex and do not require the choices involved in constructing mock communities, the obvious disadvantage of using these samples is that the correct answer is not known making it difficult to benchmark different algorithms.

In this manuscript, we consider for Kraken2 and Metaphaln2, the two most widely used classification algorithms, two traits of classification performance on real datasets beyond “correct classification” that have not previously been extensively examined in benchmarking papers. The first of these is inference. For four “real” (naturally derived communities) datasets we examine the pattern of inference and ask how many significant taxa each classifier finds. In addition, we consider the structure of Spearman correlation patterns between the reported taxa for each classifier. Despite not knowing which classification calls are “correct”, we argue that even with an overall very low error rate, given enough sequencing depth, Kraken2, but not Metaphlan2 or 16S classifiers, reliably produce many “phantom” taxa, that reflect systematic classification error, that nonetheless are significantly associated with metadata.

## Methods

The four datasets that have been chosen for this comparison (Table 1) include three human gut whole genome studies [25–27] and one pig gut whole genome classifier study [28]. All four of the datasets are available on NCBI: Vanderbilt project is PRJNA693850 (WGS) and PRJNA397450 (16S), Pig Gut study is PRJEB11755, IBD study is PRJNA389280, and the China study is PRJNA349463. Samples that did not have corresponding metadata were not included in the analysis.

**Table 1:** Metadata and Sequencing Depth at Genus Level.

| Dataset | Metadata | Sample Size WGS | Average Sequence Depth Kraken | Average Sequence Depth Metaphlan | Sample Size 16S | Average Sequence Depth RDP | Average Sequence Depth QIIME |
| --- | --- | --- | --- | --- | --- | --- | --- |
| China (Winglee et al., 2017) | Rural vs Urban | 40 | 3,668,438 | 3,463,600 | 40 | 69,021.02 | 40,830.70 |
| Vanderbilt (S. Sun et al., 2021) | Sample Type (stool vs. swab vs. tissue) | 1,203 | 2,746,762 | 2,043,168 | 240 | 134,289.20 | 107,769.70 |
| IBD (Schirmer et al., 2018) | Disease vs Control | 341 | 5,637,535 | 3,049,583 | NA | NA | NA |
| Pig Gut (Xiao et al., 2016) | 3 countries (China, France, Denmark) | 281 | 1,476,263 | 1,017,447 | NA | NA | NA |

Reverse reads were ignored and only forward reads were included. Contaminant reads were removed from the forward fastq files using kneaddata, package in bioBakery [29], which automatically calls on trimmomatic [30] for trimming and bowtie [31] to align reads to contaminant genomes. Two classifiers were tested in this study due to popularity and ease of use which were Metaphlan2, and Kraken2. These classifiers constitute the two most highly cited and used form of classification of whole genome shotgun data with Metaphlan representing marker gene based classifiers and Kraken representing k-mer based classifiers. This study used the default database for Metaphlan2. For Kraken2, we exclusively used the prokaryotic combination of refseq databases downloaded from https://benlangmead.github.io/aws-indexes/k2 that only included archaea and bacteria as downloaded on April 2020.

Statistical Analysis started with normalizing taxa tables from all classifiers through the Normalization Formula, 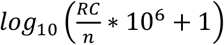, and subdividing by phylogenetic level. RC represents the raw counts value of a given taxa for a given sample, n is the total number of sequence counts for that sample, multiplying by a million normalized the proportions to the same sequence depth to allow for comparisons across different datasets, and a pseudo count of 1 was chosen to make zeros appear as zeros after log transformation. This formula is designed to minimize differences in the impact of the pseudo count and differences caused by different datasets with different sampling depths. Nonparametric Kruskal Wallis test were employed to highlight the differences in the microbiome for the variable with the strongest effect size in each dataset. In the case of the Vanderbilt dataset, this was the type of method for collecting samples either swabbing, tissue, or using patient stool. The Pig Gut dataset used the geographical location of the pig which consisted of three locations China, Denmark, and France. The IBD dataset was simply whether the patient suffered from Intestinal Bowel Disease or did not. Lastly the China dataset looked at whether the patient lived in a rural community or an urban community. All coefficient of determination and p-values reported came from these variables modeled to their datasets. FDR correction was via the Benjamini-Hochberg [32] method at a 5% FDR threshold.

Additional statistical tests consisted of calculating the highest Spearman correlation Rho values for each taxa when compared to every other taxa with higher abundance in the dataset. In this calculation, we sort each taxa based on its mean relative abundance and, starting with the 2^nd^ most abundant taxa, report the highest Spearman correlation for each taxa with higher abundance. So, for the 2^nd^ most abundant taxa, we report the Spearman correlation with the 1^st^ most abundance taxa. For the 3^rd^ most abundant taxa, we report the higher of the Spearman correlation coefficients between the 3^rd^ and the 1^st^ taxa and the 3^rd^ and the 2^nd^. And so forth until each taxa (except the most abundant) is assigned such a value. To avoid artifacts associated with normalization, Spearman correlation coefficients were calculated on un-normalized data. Performing the results on log-normalized data, however, yielded nearly identical results (data not shown).

All code, including code for the Poisson simulations described in the results is available at https://github.com/JBJohnson95/Ranking_WGS_Classifier.

## Results

### Kraken2 finds both more taxa and more taxa significantly associated with metadata than Metaphlan

In this analysis, we compared average relative abundance across all samples for four previously published publicly available datasets using Metaphlan2 and Kraken2 classifiers. In addition, for each dataset, we performed inference with relevant metadata using non-parametric tests (see methods). When comparing Metaphlan2 and Kraken2 (Figures 1–2), we noticed a strikingly similar pattern across all 4 datasets with the following features: (i) Both classifiers found taxa that were not found by the other classifier, but Kraken2 reported many more of these than Metaphlan2 at both the phyla (Fig.1) and genus (Fig. 2) level; (ii) for high abundance taxa, the two classifiers agreed very strongly on the average relative log-normalized abundances for all datasets; (iii) for all datasets there were a number of low abundance taxa which Kraken2 consistently reported a lower log-normalized relative abundance than Metaphlan2; and (iv) Kraken not only reports more taxa but more taxa that are significantly associated with metadata at both the phyla and genus level.

**Figure 1:**
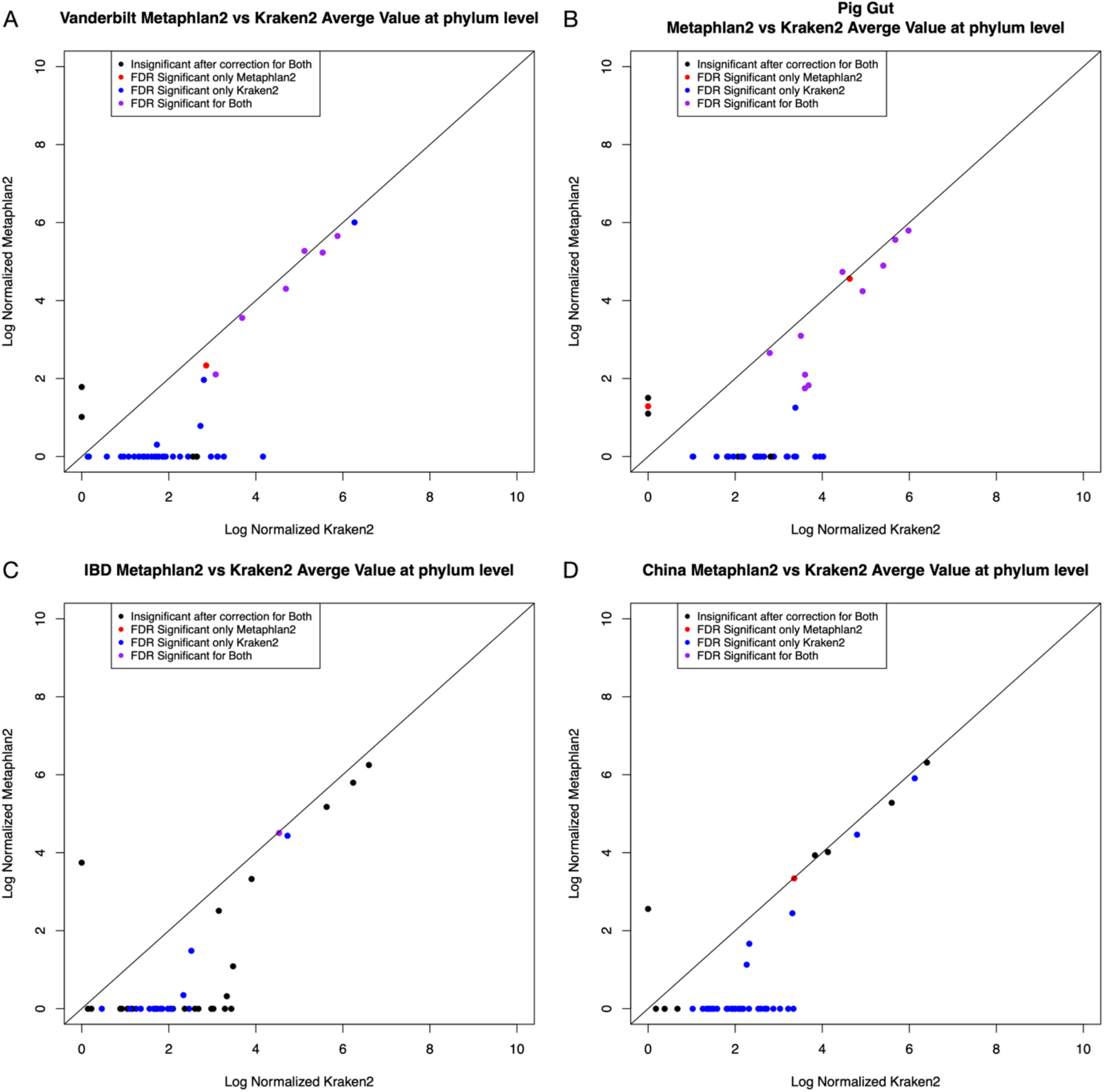
Kraken2 and Metaphlan2 normalized sample counts at the phylum level for our four publicly available datasets. Colors indicate the results of statistical tests measuring the association of each taxa with metadata for each dataset at a 5% FDR (see methods).

**Figure 2:**
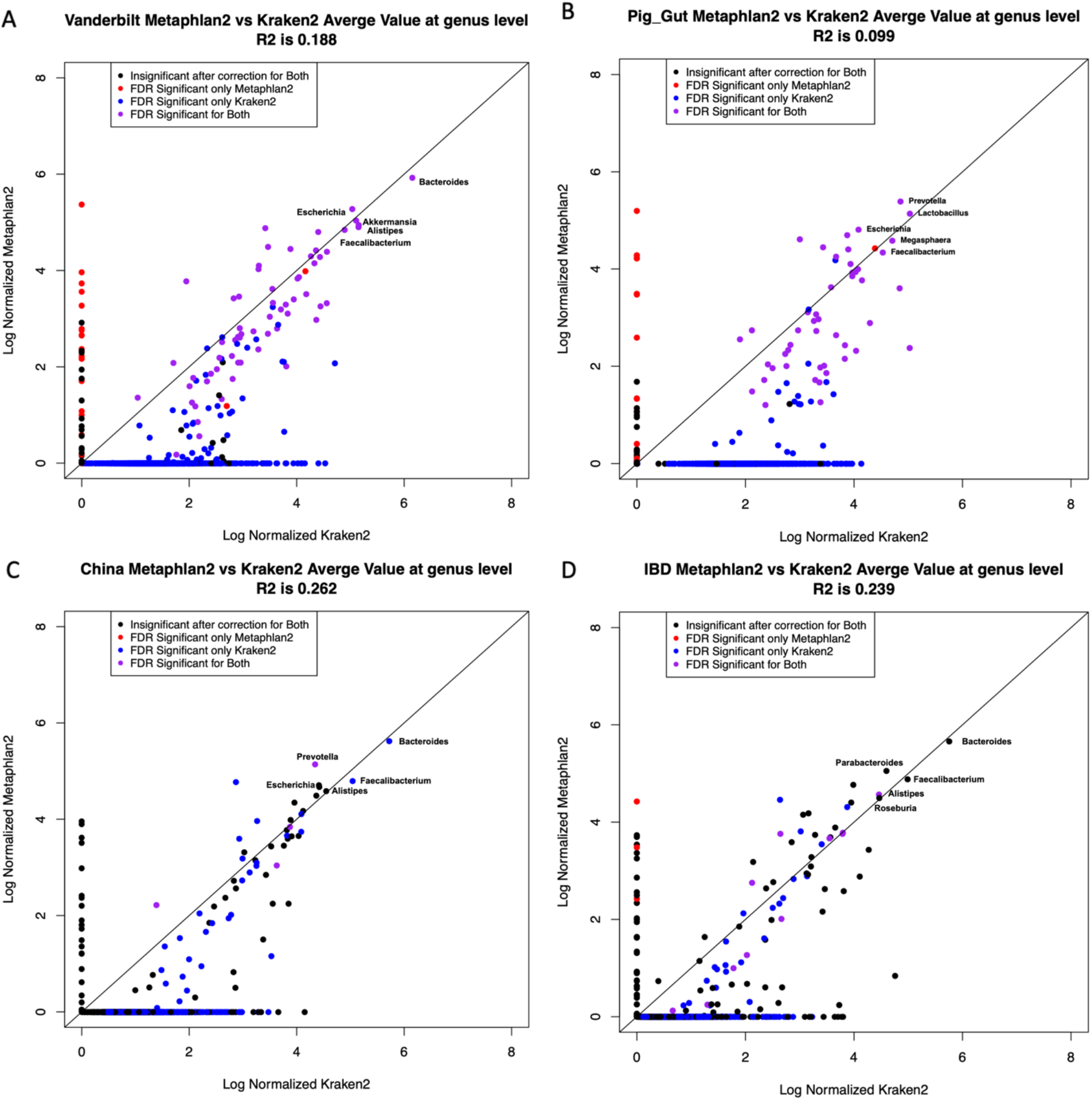
Kraken2 and Metaphlan2 normalized sample counts at the genus level for our four publicly available datasets. Colors indicate the results of statistical tests measuring the association of each taxa with metadata for each dataset at a 5% FDR (see methods).

### Patterns of Spearman correlation between taxa suggests that many classifications from Kraken2 are “phantom” taxa that represent systematic misclassification of high-abundance taxa

The fact that Kraken reports not only more taxa but also more taxa significantly associated with metadata suggests two alternative hypotheses. The first is that Kraken is more sensitive and is reporting biologically important taxa that Metaphlan simply misses. An alternative, not necessarily mutually exclusive hypothesis, is that Kraken is misclassifying some reads in a non-random way that impacts inference. In order to discriminate between these two hypotheses, we asked for each taxa what was the highest Spearman correlation coefficient with any more abundant taxa (see methods). Surprisingly we discovered that the vast majority of the taxa that Kraken2 identified appeared to be highly correlated with at least one more abundant taxa and therefore shared that taxa’s distribution across samples (Fig. 3–6). Metaphlan2 did not show a similar correlation structure (Figs. 3–6).

**Figure 3:**
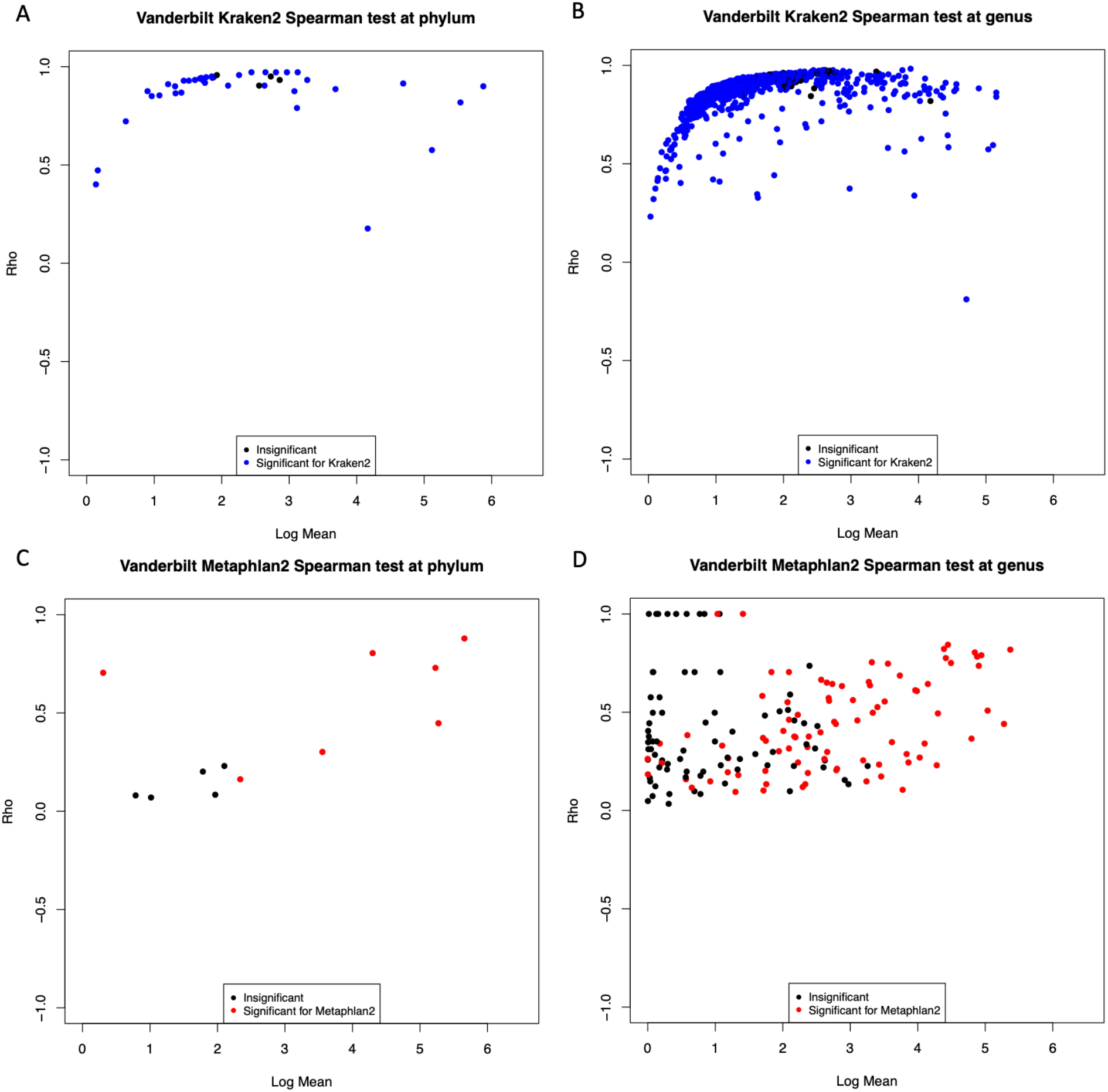
For each taxa, the highest Spearman correlation coefficient (y-axis) with any other higher abundance taxa in the Vanderbilt dataset. The x-axis shows the normalized log-abundance for each taxa. Colors indicate the results of statistical tests measuring the association of each taxa with differences in testing site for the Vanderbilt dataset at a 5% FDR (see methods).

**Figure 4:**
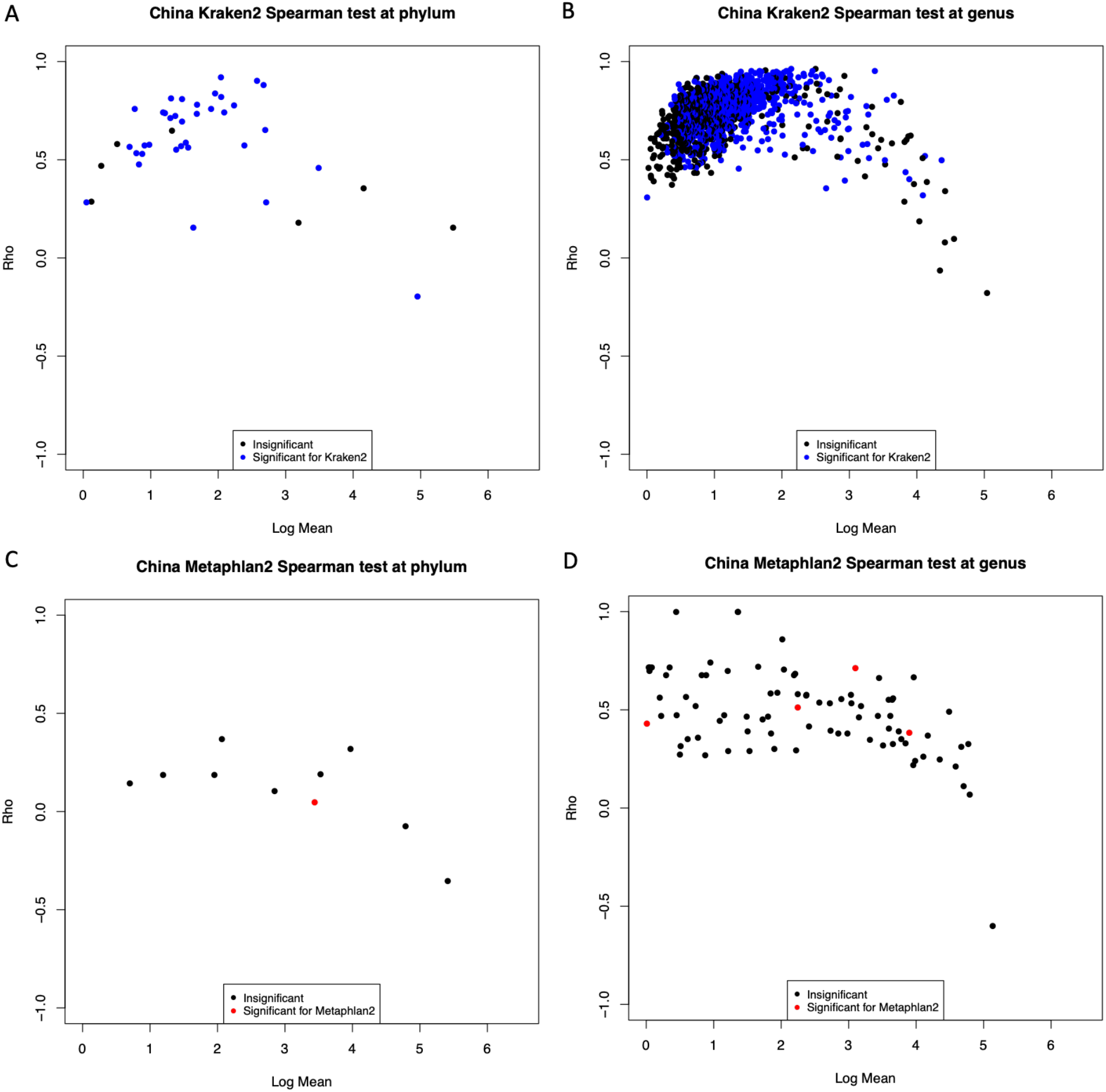
For each taxa, the highest Spearman correlation coefficient (y-axis) with any other higher abundance taxa in the China dataset. The x-axis shows the normalized log-abundance for each taxa. Colors indicate the results of statistical tests measuring the association of each taxa with differences in geographical urban density reduced to a binary choice of urban or rural for the China dataset (see methods).

**Figure 5:**
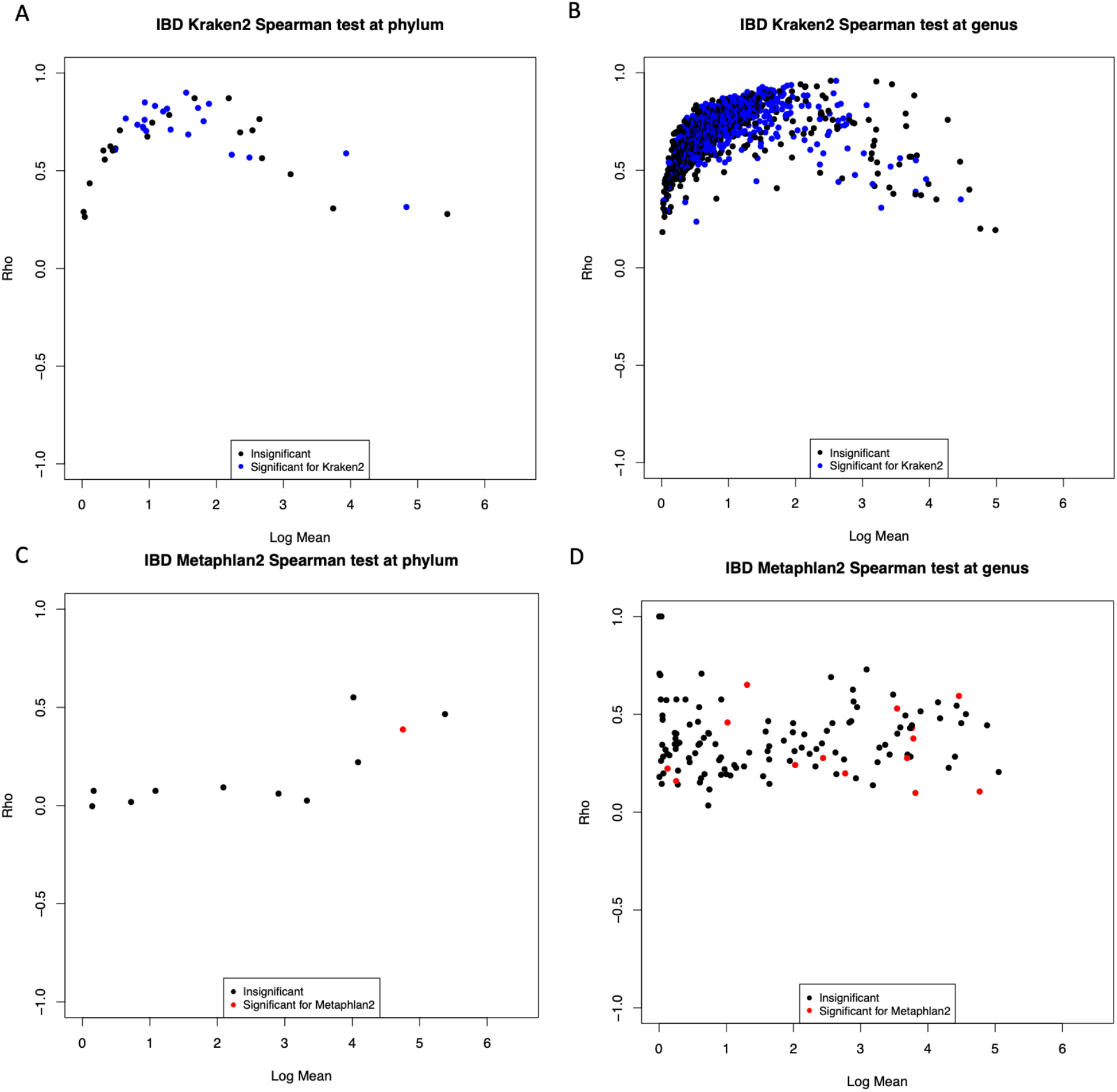
For each taxa, the highest Spearman correlation coefficient (y-axis) with any other higher abundance taxa in the IBD dataset. The x-axis shows the normalized log-abundance for each taxa. Colors indicate the results of statistical tests measuring the association of each taxa with differences in case/control for IBD (see methods).

**Figure 6:**
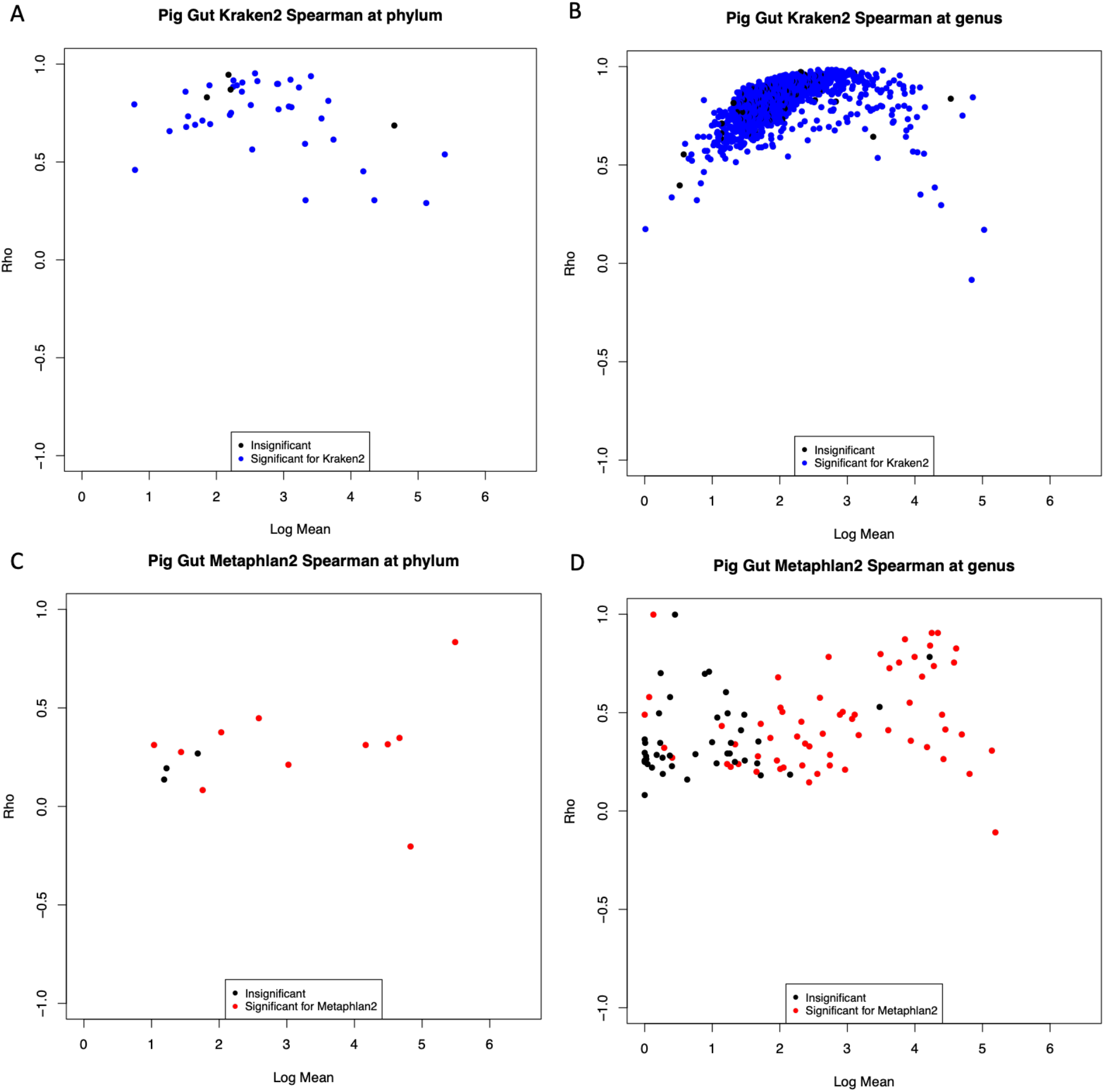
For each taxa, the highest Spearman correlation coefficient (y-axis) with any other higher abundance taxa in the Pig Gut dataset. The x-axis shows the normalized log-abundance for each taxa. Colors indicate the results of statistical tests measuring the association of each taxa with differences in geographical location of the pig gut microbiomes observed for the Pig Gut dataset (see methods).

One possible explanation for this observation is that there exists in these samples many low abundance taxa that are truly correlated with the high abundance taxa that Kraken is able to classify but that are ignored by the more conservative Metaphlan2. To explore this possibility, we examined matched 16S sequences which were available for two of our four datasets. We classified the 16S sequences with RDP and QIIME and found that like WGS classifiers, the two 16S classifiers showed reasonable concordance for high abundance taxa but distinct differences for many lower abundance taxa (Fig. 7). However, when we calculated for each taxa the maximum Spearman correlation coefficient with another higher abundance taxa, we did not observe the high degree of correlations that we found for Kraken (Fig. 8–9) despite the fact that Kraken2 and Metaphlan2 tend to agree with both 16S classifiers for the high abundance taxa (Fig. 10–11) with Kraken2 again having many more lower-abundance taxa detected than Metaphlan2 or either of the 16S classifiers. If the highly correlated low-abundance taxa detected by Kraken were truly present, it is reasonable to assert that the correlation structure would be detected by both WGS and 16S. The fact that we did not observe this leads us to argue that the low abundance taxa detected by Kraken are better explained as classification error.

**Figure 7:**
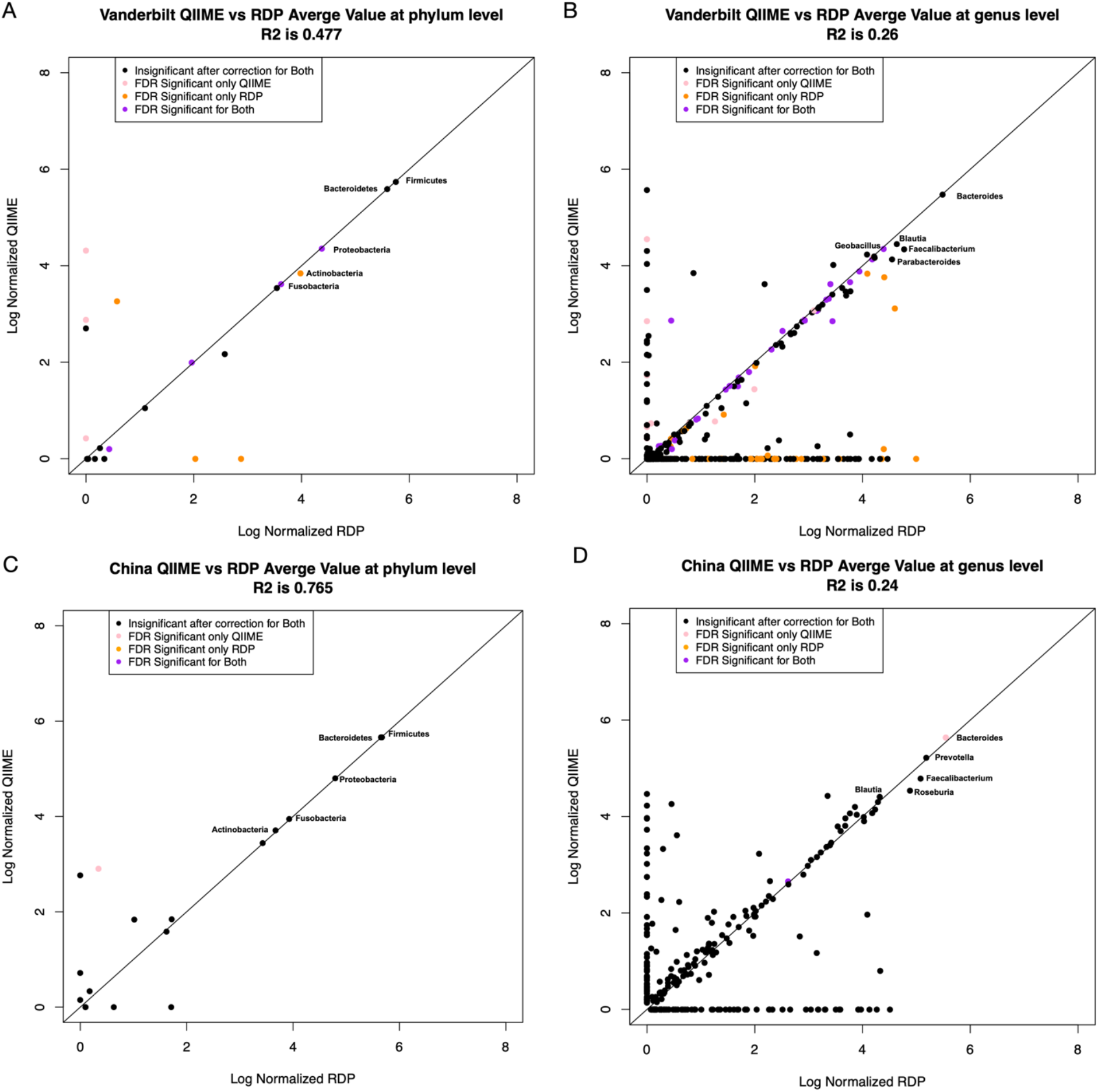
Normalized sample counts at the phylum and genus level were averaged by taxa for both 16S classifiers RDP on the x-axis and QIIME on the y-axis. Phylum level is on the left column and Genus level is on the right column. A) Vanderbilt Phylum B) Vanderbilt Genus C) China Phylum D) China genus.

**Figure 8:**
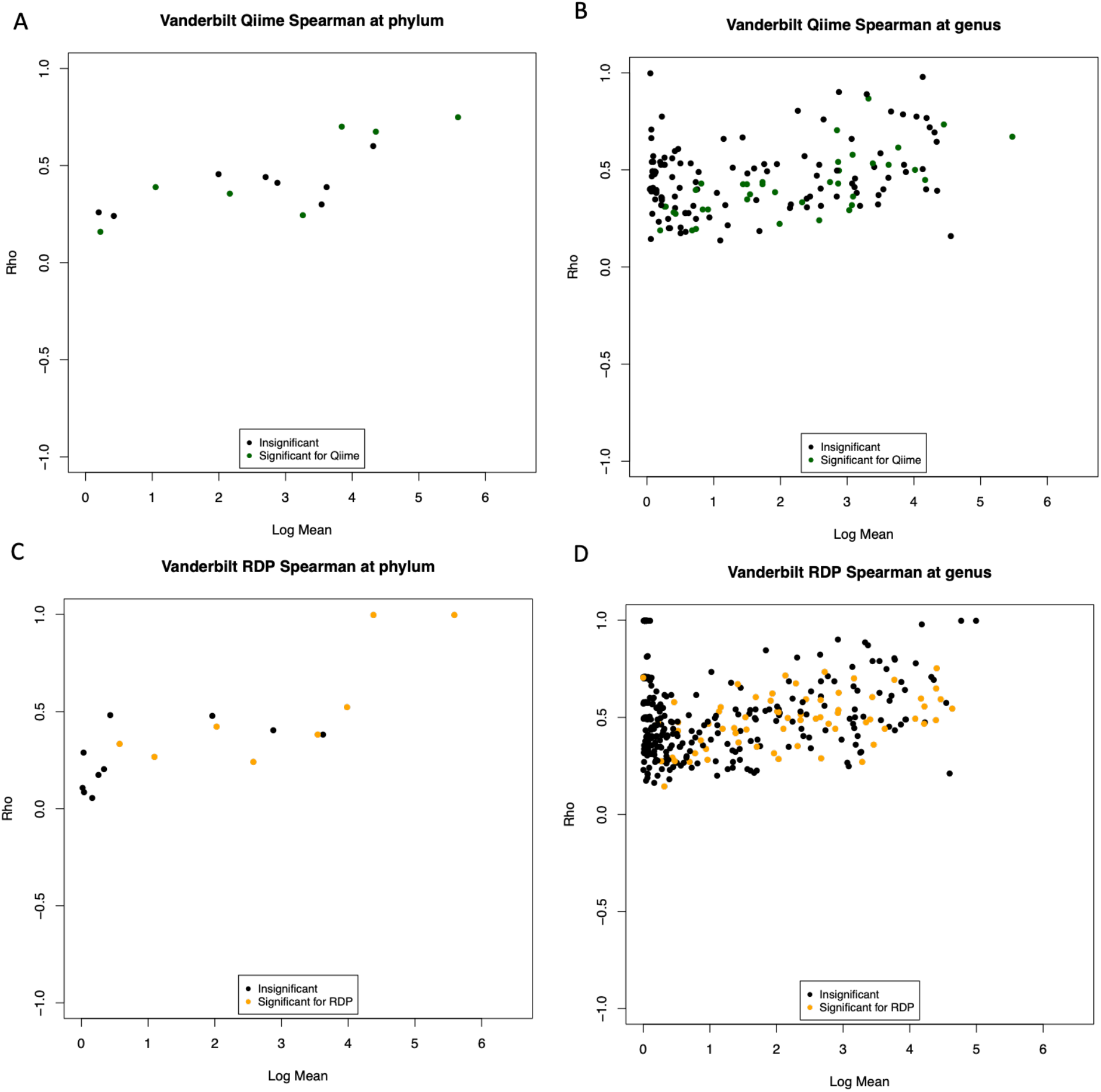
Spearman Correlations (the highest correlation coefficient (y-axis) with any other taxa in each dataset) for QIIME and RDP for Vanderbilt dataset plotted against the log average mean for each taxa. Figures (A-B) are for QIIME and Figures (C-D) are for RDP. Phylum level is on the left column (A and C) and Genus level is on the right column (B and D).

**Figure 9:**
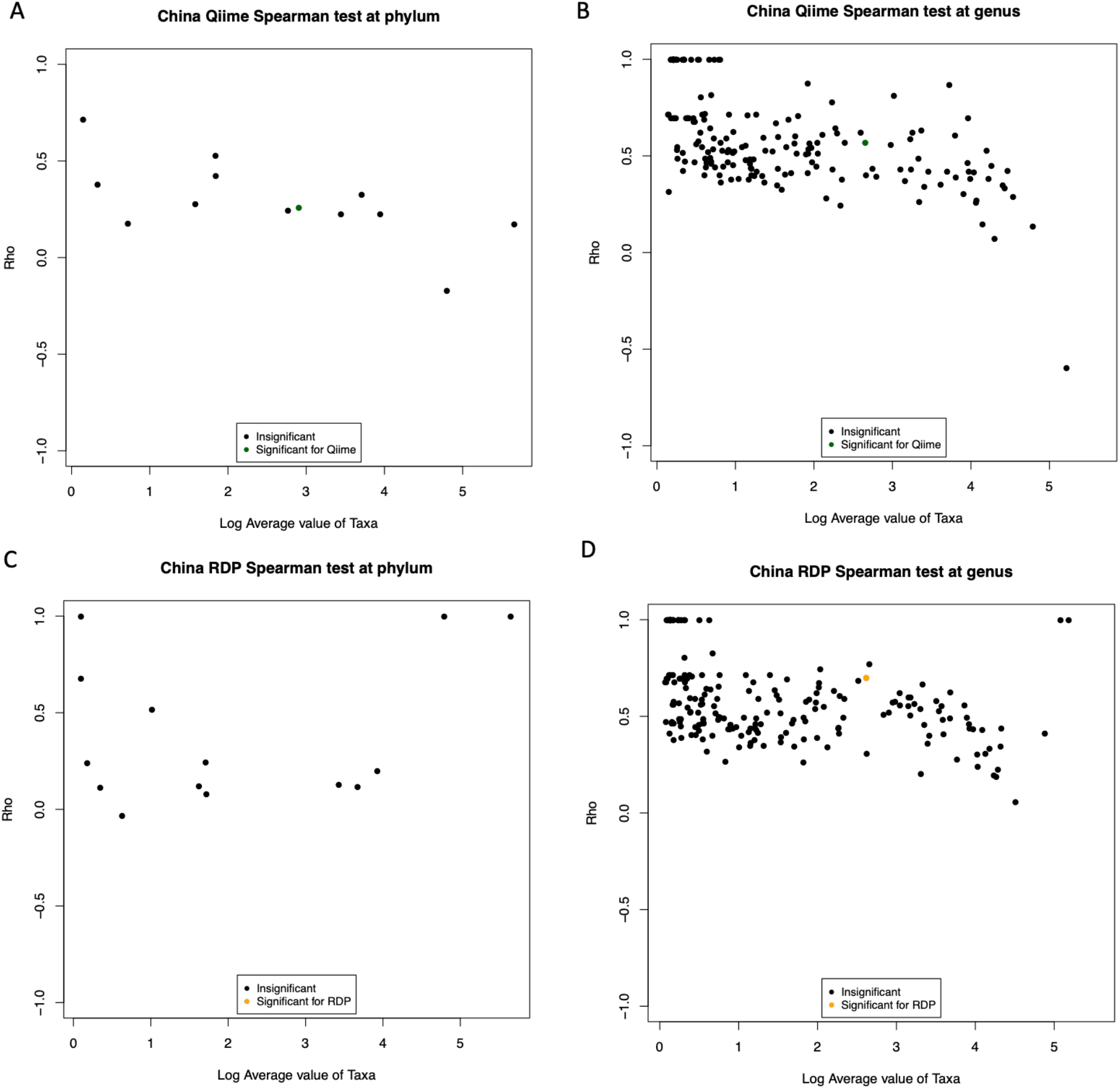
Spearman Correlations (the highest correlation coefficient (y-axis) with any other taxa in each dataset) for QIIME and RDP for China dataset plotted against the log average mean for each taxa. Figures (A-B) are for QIIME and Figures (C-D) are for RDP. Phylum level is on the left column (A and C) and Genus level is on the right column (B and D).

**Figure 10:**
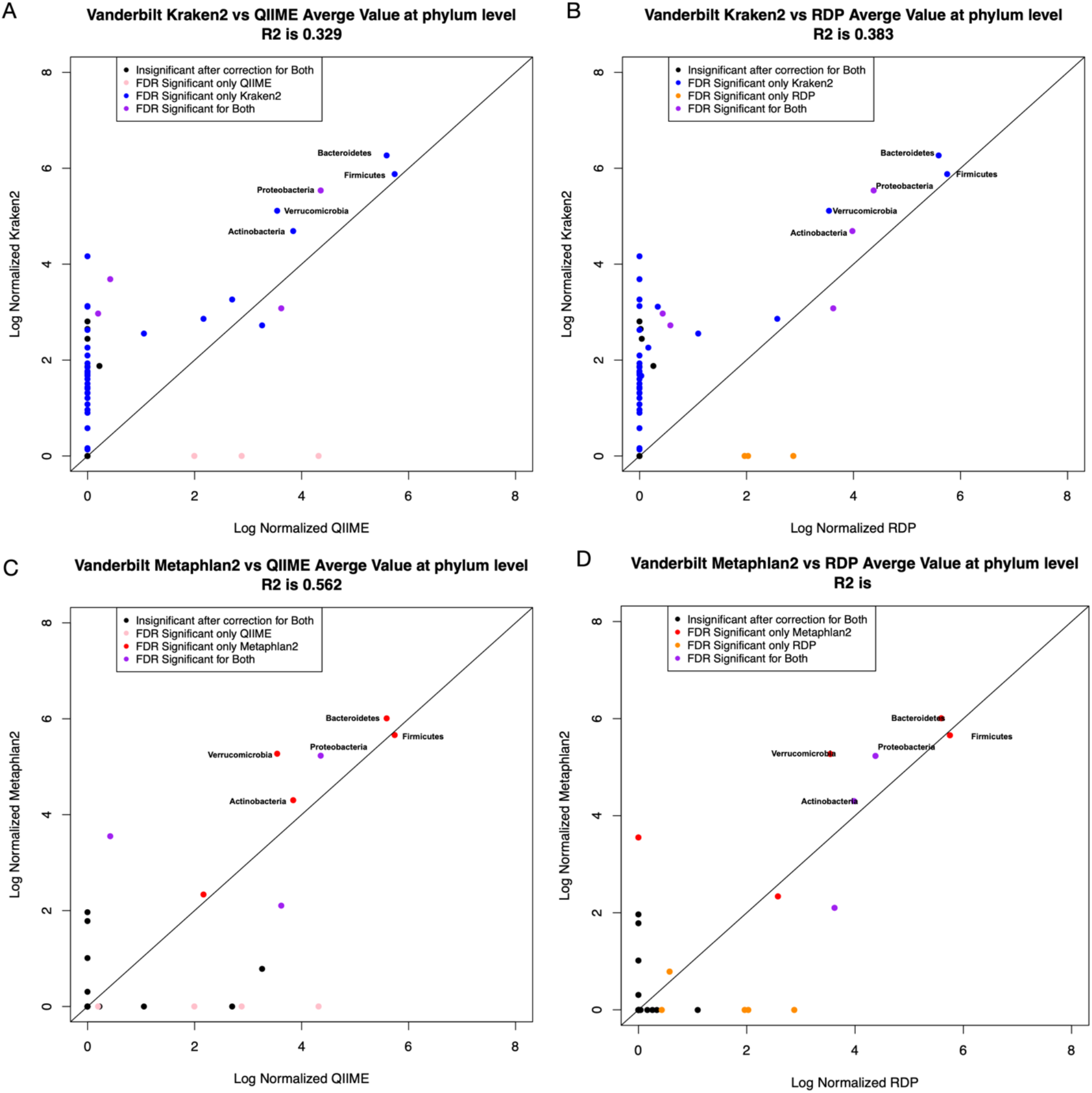
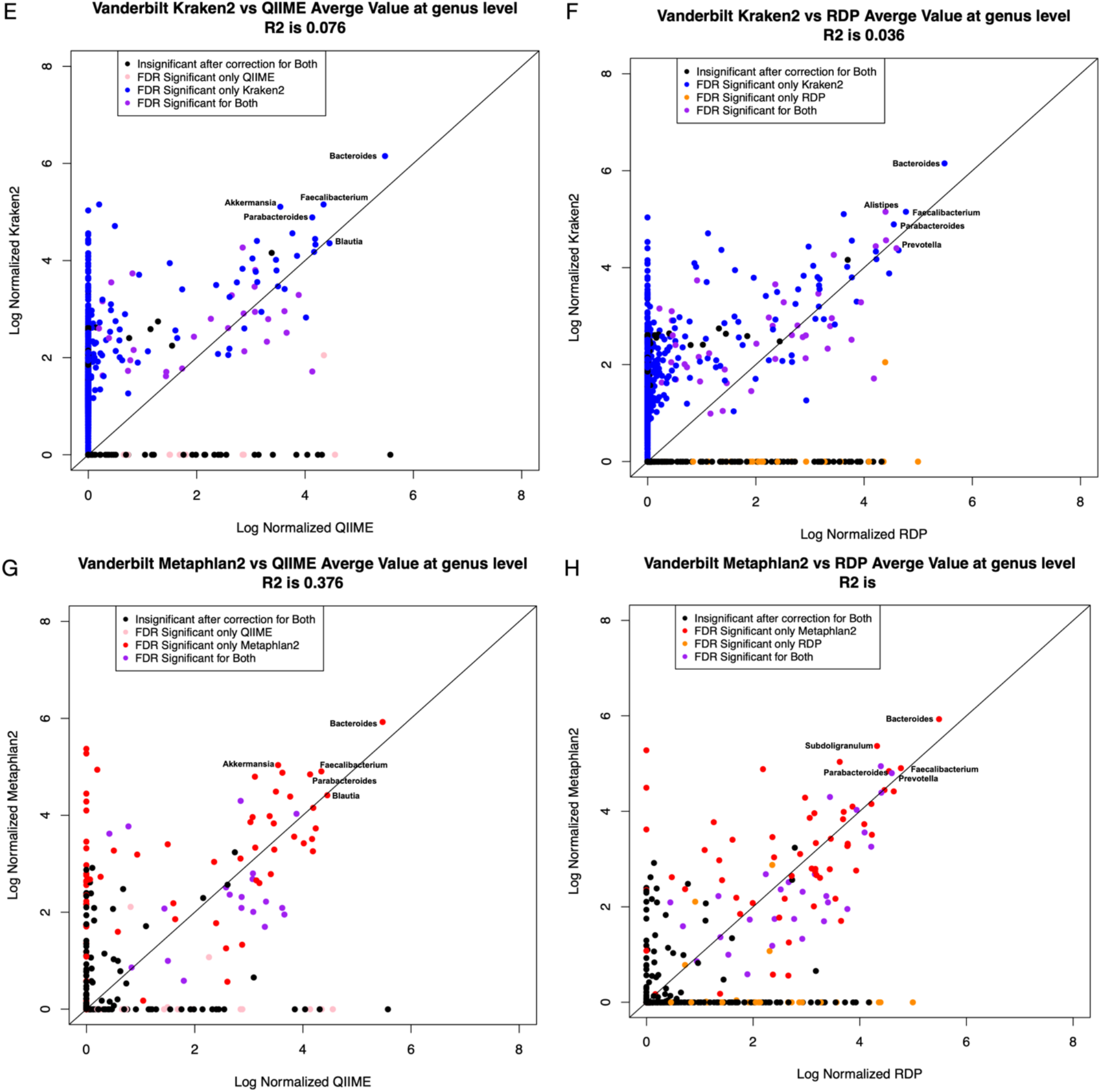
Normalized sample counts at the phylum and genus level were averaged by taxa for both 16S classifiers and WGS Classifiers in the Vanderbilt dataset. Figures A-D are at the phylum level while Figures E-H are at the genus level. Figures A-B and Figures E-F comparing Kraken2 against QIIME (A and E) and RDP (B and F) while Figures C-D and Figures G-H compare Metaphlan2 to QIIME (C and G) and RDP (D and H).

**Figure 11:**
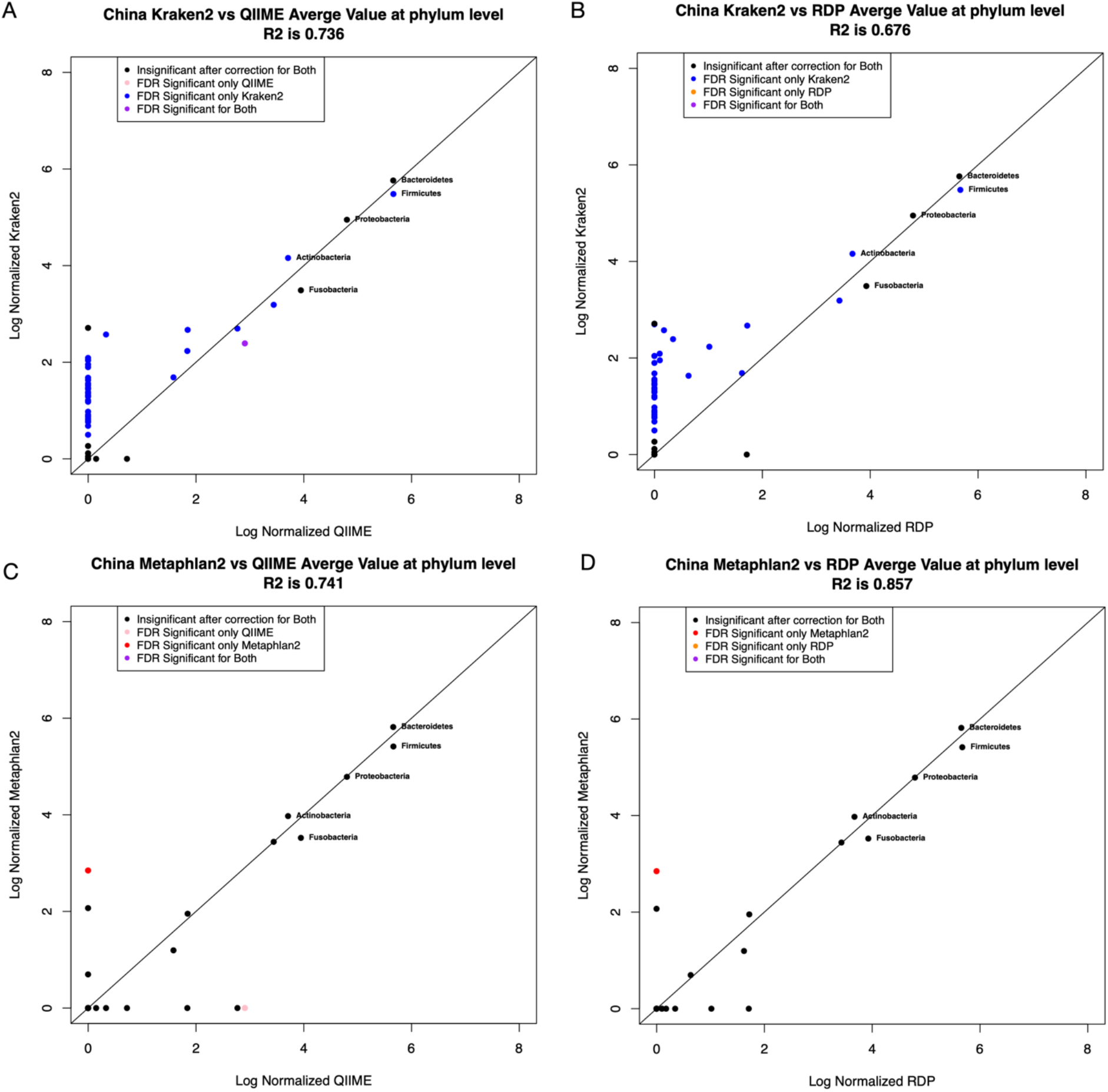
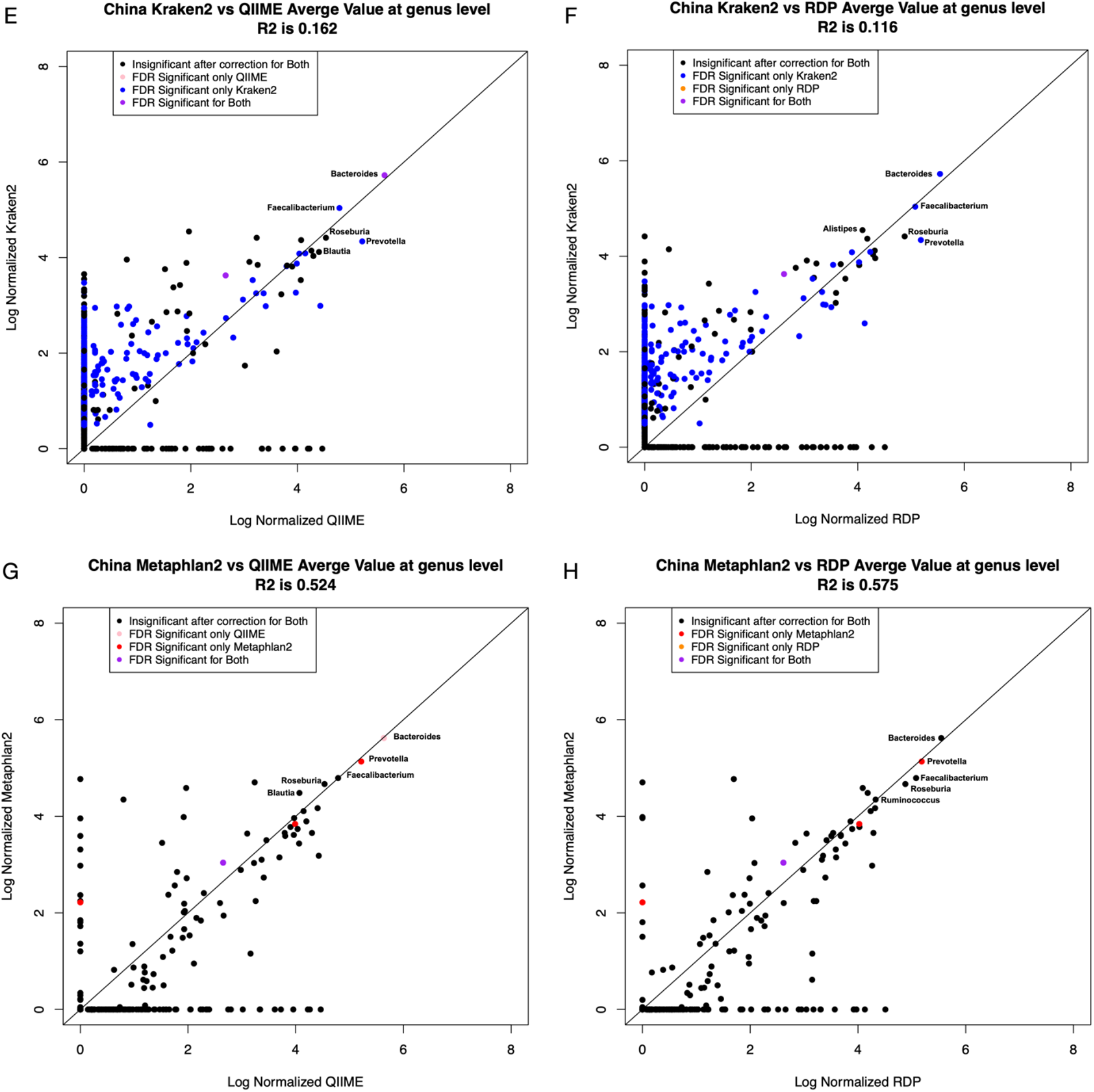
Normalized sample counts at the phylum and genus level were averaged by taxa for both 16S classifiers and WGS Classifiers in the China dataset. Figures A-D are at the phylum level while Figures E-H are at the genus level. Figures A-B and Figures E-F comparing Kraken2 against QIIME (A and E) and RDP (B and F) while Figures C-D and Figures G-H compare Metaphlan2 to QIIME (C and G) and RDP (D and H).

### A very simple Poisson model of classification error with a low error rate sampled from the uniform distribution shows good concordance with Kraken’s correlation structure

In order to test the idea that “phantom” taxa could be generated from very simple models of mis-classification, we created a simple Poisson based model. In this model, we assumed that the 10 most abundant taxa in a sample are “real”. Then for each taxa not in the top ten, we estimate an abundance in each sample with an assumption that the abundance was only caused by classification error. For this process, we choose one of the 10 most abundant taxa in a weighted way. For example, if the most abundant taxa represented 50% of all the reads of the 10 most abundant taxa, we would choose those taxa 50% of the time. If the next most abundant taxa were present 30% of the time, we would choose those taxa 30% of the time and so forth. We then assume an error rate that is different for each “phantom” taxa sampled from the uniform distribution from 0 to 0.001 (for an average error rate of 1 in 2000 reads). For each taxa in each sample, we use the Poisson distribution with lambda set to the number of reads of the chosen “real” taxa in that sample times the error rate sampled from the uniform distribution. In this way we generate a counts table in which the top 10 taxa are what was observed from the sequencer and all the other taxa represent classification error. As can be seen in Figures 12–13, for Kraken the correlation structure of these tables in which we graph the correlation coefficient of each “phantom” taxa to its “real” parent (green dots in Figs. 12–13) is a very reasonable match to the actual correlation structure where we graph the maximum correlation coefficient of each taxa to all other taxa in the sample (non-green dots in Figs. 12–13). This model, however, fails to match the correlation patterns of either Metaphlan or either 16S classifier.

**Figure 12:**
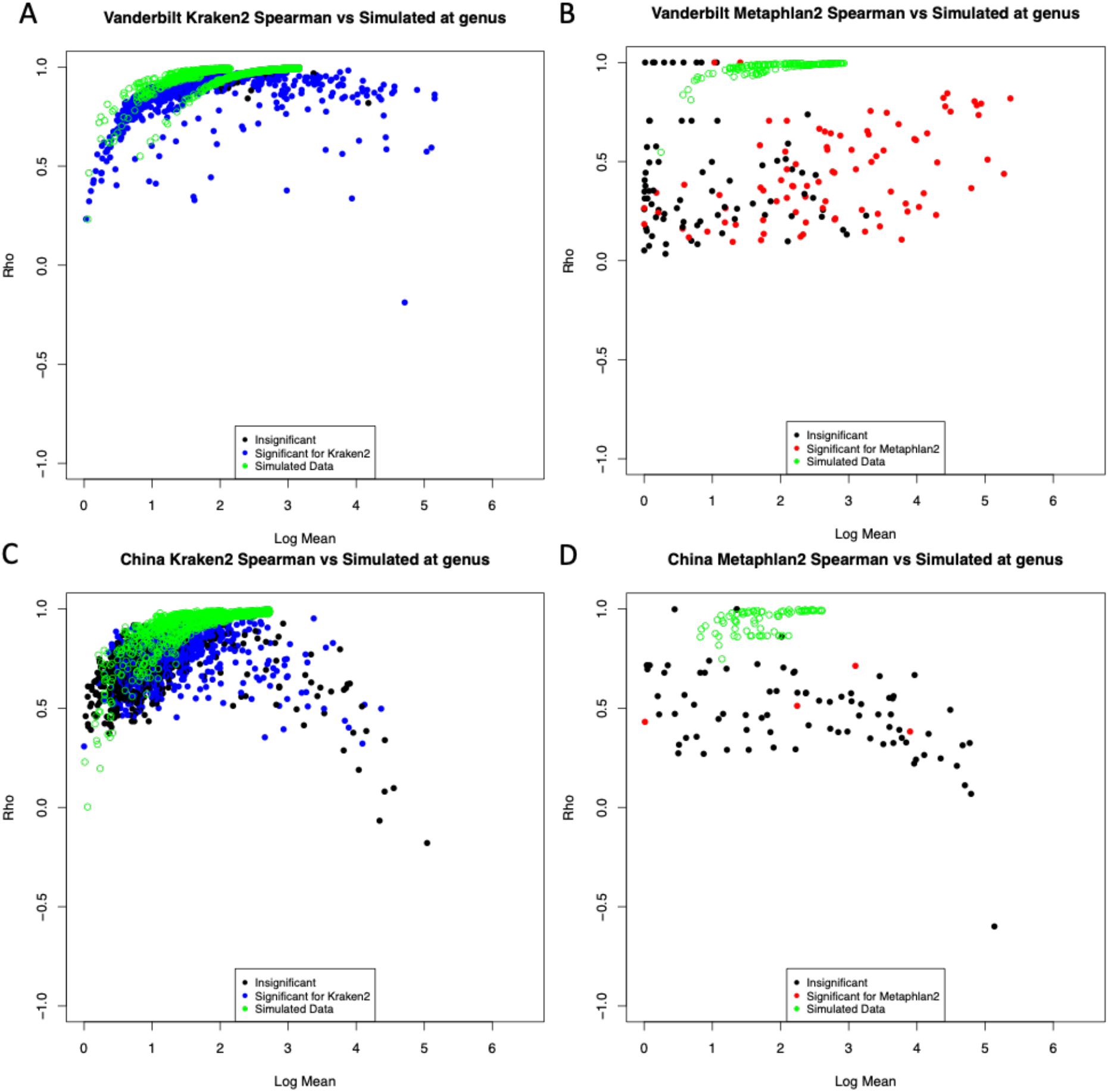
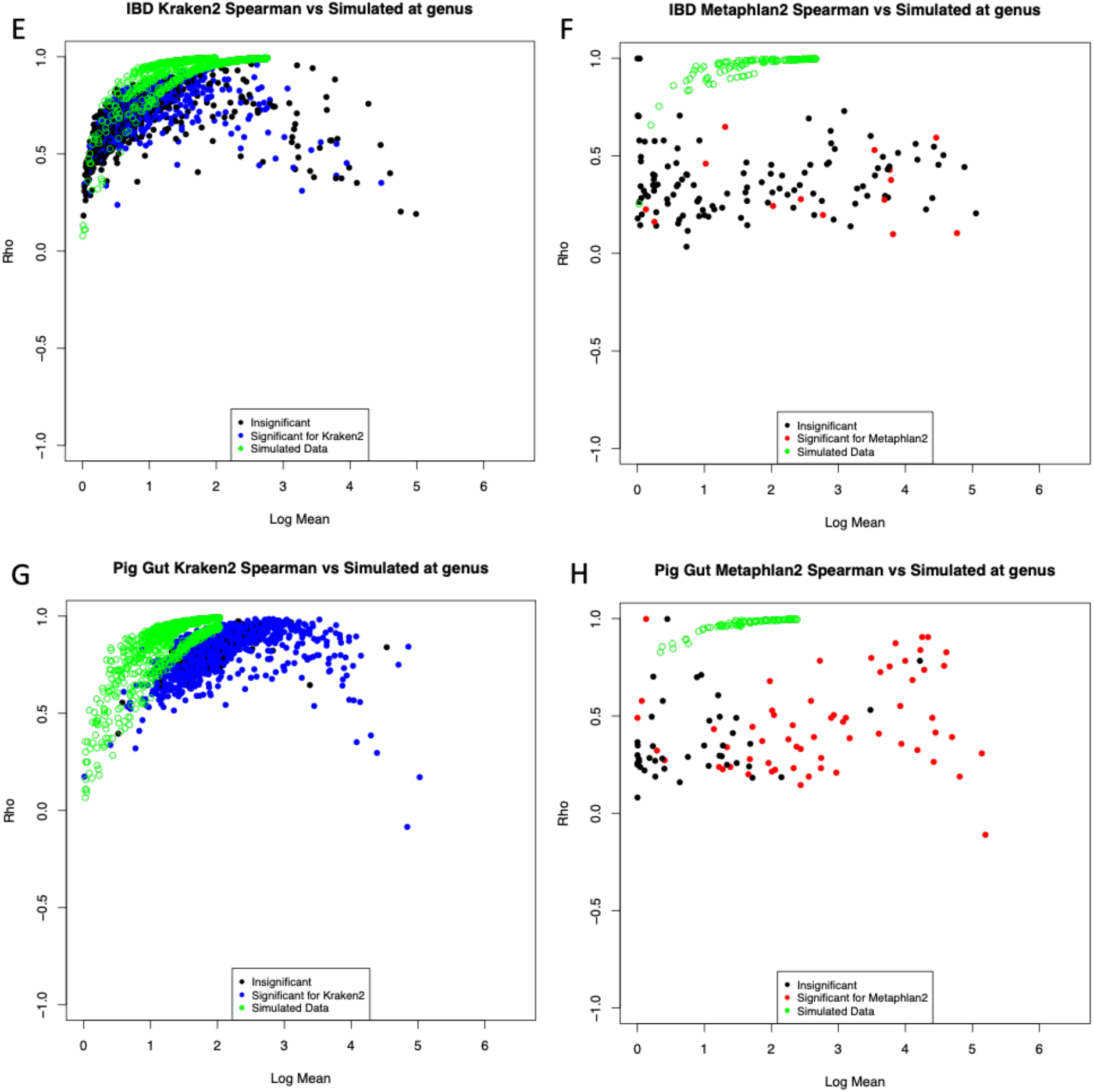
Normalized sample counts averaged by taxa at genus level were compared against a simulated binomial distribution of phantom taxa labeled in green. Kraken2 results are shown on the left column (A,C,E,G) while Metaphlan2 results are shown on the right column (B,D,F,H). The y-axis is the highest Spearman correlation against each other high abundance taxa.

**Figure 13:**
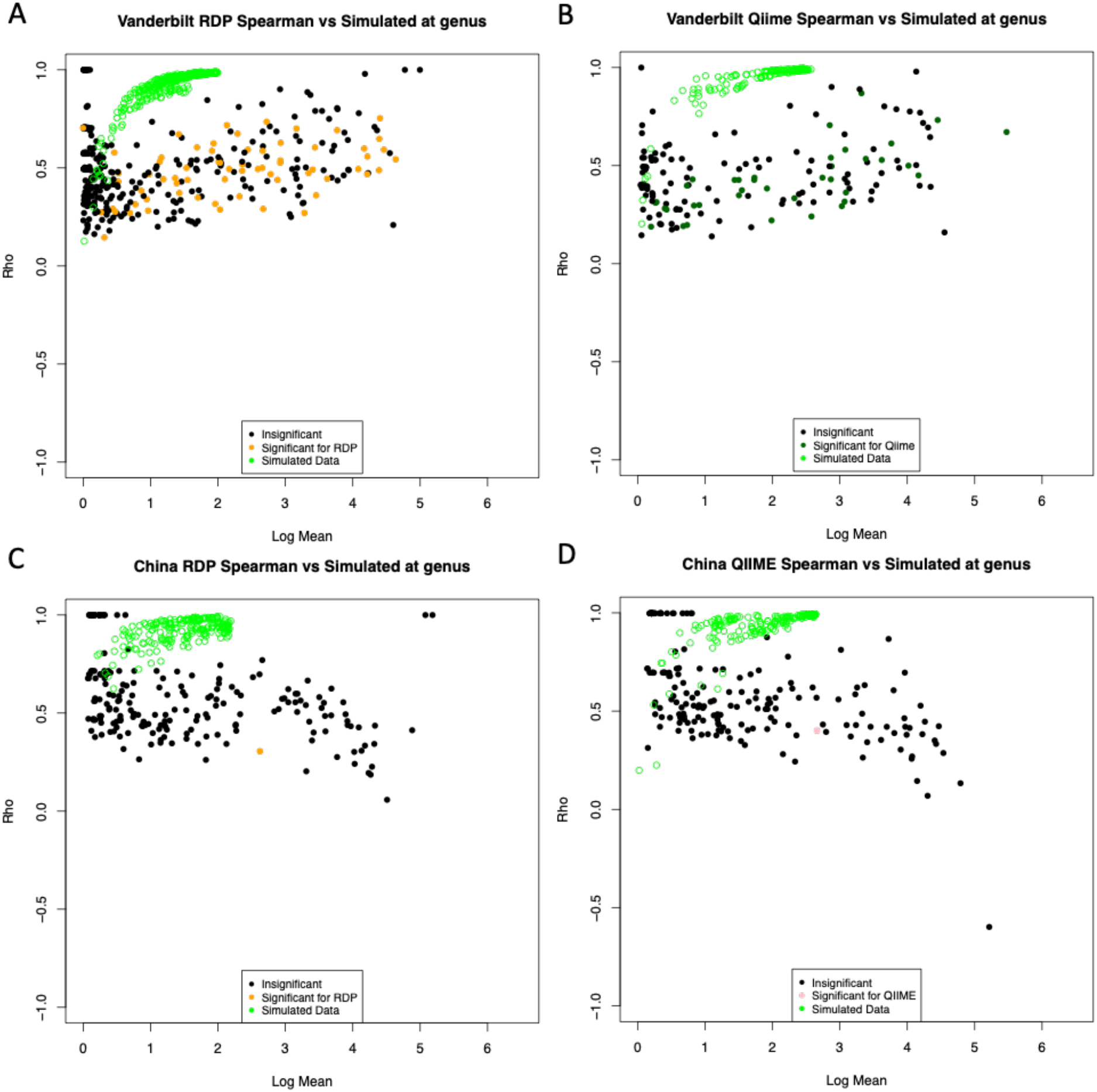
16S normalized sample counts averaged by taxa at the genus level were compared against a simulated binomial distribution of phantom taxa labeled in green. RDP results are shown in the left column (A,C) and QIIME results are shown in the right column (B,D). The y-axis is the highest Spearman correlation against each other high abundance taxa.

We conclude that the behavior of Kraken, but not Metaphlan or 16S classifiers on 16S data, is consistent with a pattern of classification error where some fraction of high abundance taxa are systematically misclassified as lower abundance taxa.

## Discussion

Our analysis has immediate implications for biologists who wish to use taxonomic patterns for inference based on WGS data. For the most highly abundant taxa, use of Kraken2 or Metaphlan2, or for that matter analysis of 16S sequences with QIIME or RDP, will yield highly similar answers. If there are no high abundance taxa that are significantly associated with metadata variables of interest, then likewise the choice of sequencing technology or algorithm will likely have a limited impact on inference. However, as is often the case in biological datasets and appears to be the case for the four datasets that we have chosen, if there is a high abundance taxa that is associated with metadata, and a sufficient sequencing depth, there is a possibility that Kraken2, but not Metaphlan2 or 16S classifiers, will have a small but systematic classification error in which members of the high abundance taxa are mis-characterized as a different taxa. Evidence for this assertion can be seen in our observation that Kraken, but not Metaphlan or 16S classifiers, routinely produces many taxa that have very high correlation coefficients with more abundant taxa. This leads to the danger that these “phantom” taxa, that share a distribution with a “real” more abundant taxon that is correlated with metadata, will produce a spurious significant result in inference leading to the conclusion of a significant association with a taxon that is either not present in the samples or present but with a distribution that has been distorted by a mix of true reads and misclassifications from a higher abundance taxa.

Supporting this idea, our very simple model with Poisson based classification error of the most abundant taxa was able to capture much of the pattern of the correlation structure of Kraken2, but not Metaphlan2 or the 16S classifiers. Our model shows that even with a very low average classification error (an average error rate of 1 read in 2,000), this type of behavior can produce many spurious correlations given adequate sequencing depth. It is somewhat remarkable that our model with sampling from the uniform distribution with a single parameter (the average error rate of 0.001) was able to describe the pattern of correlation for all the datasets so well with only the Pig Gut dataset at the genus level suggesting our model has underestimated the error rate. These results suggest an overall fairly consistent pattern of error across datasets collected from host-associated microbiomes under very different biological circumstances. It would be of interest in future work to see if environmental datasets such as soil or ocean, which we did not consider in the current work, had an overall higher error rate when viewed through this prism of correlation structure. Our results suggest that benchmarking studies should consider not only absolute error rate but patterns of how misclassifications occur. A classifier that randomly misclassifies a read to one of a large number of taxa with a higher overall error rate will not generate this pattern as much as a classifier that systematically misclassifies reads even with a much lower error rate.

Ironically, this problem is made more serious by the very high sequencing depths made possible by technological advancements in the last few years. As sequencing depth has increased recently to the range of billions of reads per sample, even a relatively low error rate can produce quantifiable problems in terms of classification, as our Poisson modeling shows. As a hypothetical example, consider an error rate of 1% on a taxon with a relative abundance of 10%. For a dataset with 100,000 reads per sample, the taxon would have on average 10,000 reads per sample and mis-classification would only produce 100 reads, which might not be enough to survive filters for low abundance taxa or successfully perform inference across samples. However, with modern technology, it is easy to achieve 100 million reads per sample. At this sequencing depth, the taxa would produce 10 million reads and a 1% error rate would produce 100,000 reads devoted to a “phantom” taxa, which of course will be highly correlated across samples with the “real” highly abundant taxa so that we would expect a common pattern of inference between the phantom and the real taxa. The dependence of these problems on sequencing depths means that the patterns that we report here may not have been observable to previous evaluation papers that used older sequence datasets with less sequencing depth. We note as a limitation to our analysis that the 16S sequencing depths in our datasets are much lower than the WGS sequencing depths (Table 1). This may explain why we fail to see the same pattern of correlation coefficients in the 16S datasets that we saw for Kraken and would require datasets with more varied sequencing depth to properly test. We also note, that these patterns of correlation structure would not be detectable in typical MDS or PCoA analysis as all of the highly correlated taxa would be well represented by individual principle components or coordinates. However, alpha diversity estimates, especially richness, have the potential to be profoundly impacted by this sort of systematic classification error.

A recent benchmarking paper [24], argues that due to the difference in taxonomic abundance vs sequence abundance researchers should proceed with extreme caution in comparing classifiers such as Kraken with algorithms such as Metaphlan. However, we find that for high abundance taxa there is very strong agreement between Kraken and Metaphlan despite the differences between the normalization schemes. We argue based on the differences in correlation structure that differences between Metaphlan and Kraken are in part explained by differences in patterns of classification error in addition to the differences in normalization. Our results suggest that caution must especially be used in construction of network diagrams that explicitly depend on correlation coefficient. The fragility of these correlation coefficients in relative abundance data has been well studied and many algorithms attempt to correct for compositional artifacts [33, 34]. We note that correlations produced by systematic classification error of high abundance taxa are another source of potential error. We anticipate development of algorithms that can potentially correct for these sources of error as a priority for bioinformatics especially as sequencing depths continue to improve with development of new technology.

While our work does suggest that these problems are more serious for Kraken vs. Metaphlan, it remains an open question whether Metaphlan’s curated marker database approach is too conservative and may be missing some true taxa that Kraken is able to be classify. Our work suggests that further development in the area of WGS classifiers is warranted and that there are many unsolved problems in this area. Given this, we argue that 16S sequencing still has some utility, especially when paired with Whole Genome Shotgun Sequencing as it allows for informative comparisons with WGS classifiers. As sequencing costs continue to decline, future studies may be able to perform high-depth 16S sequencing as a control paired with high-depth WGS. Such paired studies could provide confidence that observed correlation between taxa was not the results of systematic classification error.

